# Early life pain alters the response to an immune challenge in adult male and female rats

**DOI:** 10.1101/2023.11.17.567599

**Authors:** Morgan G. Gomez, Melani Macik, Hannah J. Harder, Malika H. Pandit, Anne Z. Murphy

## Abstract

Infants born prematurely are more likely to be admitted to the Neonatal Intensive Care Unit (NICU) where they experience upwards of 10-18 painful procedures each day, often with no anesthesia or analgesia. Pre-clinical studies have shown that early exposure to pain disrupts CNS development in multiple ways that persist into adulthood, with similar findings observed clinically. The present study explores the effects of neonatal injury on the response to an immune challenge in adulthood. Male and female rats were exposed to a short-term inflammatory insult induced by intraplantar administration of 1% carrageenan (CGN) on the day of birth (P0). In adulthood (P60-P90), rats were implanted with Thermicron iButtons to monitor core body temperature; 14 days later, lipopolysaccharide (LPS) was administered to elicit an immune response. Rats were sacrificed after 24 hours or at their peak fever point and brain tissue collected for immunohistochemical analysis. LPS administration resulted in a significantly greater febrile response in males and females exposed to early life pain compared to controls. Immunohistological analysis revealed sex and treatment differences in Fos labeling in several brain regions, including the PVN. No sex or treatment differences were observed for VGat, VGlut2, or EP3R expression in the MnPO; however, all three increased significantly following LPS administration. Together, these studies are consistent with clinical studies reporting children experiencing unresolved pain during the perinatal period show an increased severity of sickness behavior and altered immune signaling following exposure to a pathogen and will provide a foundation for future studies examining the biological underpinnings.

## Introduction

Each year, 10% of infants born in the US and up to 18% worldwide are born prior to 37 weeks and are considered premature (Osterman et al., 2021; WHO, 2012). Premature infants spend an average of 25 days in the Neonatal Intensive Care Unit (NICU) where they experience upwards of 15 painful procedures each day, including heel lances, arterial and venous line placement, intubation, and surgery. The majority of these procedures are performed without the benefit of analgesia or anesthesia (Carbajal, 2008; Roofthooft et al., 2014) as historically, premature infants were thought to have immature sensory systems and thus unable to process pain (Schwaller & Fitzgerald, 2014; Rodkey & Pillai, 2013). This long-held belief persisted until the late 1980s before being challenged and is a large part of why analgesia and anesthesia are not used in most neonate procedures to this day (Rodkey & Pillai, 2013; Simons et al., 2003).

Unresolved pain during the early neonatal period is known to disrupt the normal development of the hypothalamic-pituitary-adrenal (HPA) axis (Victoria et al., 2013a; Victoria et al., 2013b). In particular, we have previously reported in rats that inflammatory pain on the day of birth results in a blunted response to acute pain and stress, and a hyper-response to chronic pain and stress in adulthood. Site specific changes in CR1, CR2, and GR have also been reported, as well as elevated corticosterone levels 7 days post injury (LaPrairie and Murphy, 2007; Victoria et al., 2013a; Victoria et al., 2013b). Glucocorticoids, a key player in the HPA axis, has known impacts on immune related genes. For example, preterm infants exposed to unresolved pain show genetic variation in the NFKBIA gene promoter resulting in the dysregulation of the nuclear transcription factor NF-κB which plays an important role in proinflammatory cytokine and chemokine production(Grunau et al., 2013).

Preterm birth interrupts the transfer of maternal antibodies, most of which occurs in the later part of the third trimester. Clinical studies report that premature infants not only have lower antibody titers at birth compared to term controls but also the rate of decrease in maternal antibodies associated with normal development is expedited in preterm infants, further increasing their vulnerability to pathogen exposure (Melville & Moss, 2013). The maternal transfer of antibodies is critical in supporting an infant’s innate immune system during the first few months of life as the adaptive immune system begins to develop. Without it, premature infants are less able to mount a proper response to a pathogen than their term counterparts (Strunk et al., 2011). These changes also influence the development of adaptive immunity such that children born preterm have lower levels of circulating lymphocytes which persists at 7 months and 8 years compared to age matched children born full term (Berrington et al., 2005; Pelkonen et al., 1999).

In response to a skin breaking injury, peripheral immune cells infiltrate the site of injury and release proinflammatory cytokines and other inflammatory molecules including prostaglandins and eicosanoids into the bloodstream (Zouikr et al., 2016; Duorson et al., 2021). Exposure to a pathogen results in a similar release of proinflammatory cytokines (e.g. IL-1β, Tnf-α, IL-6) and prostaglandin (PGE) into the circulatory system to produce a systemic response. Cytokines cross the blood brain barrier and act directly on multiple brain regions to initiate fever and sickness-associated behaviors. One such region, the hypothalamic median preoptic nucleus (MnPO), is critical for the initiation of a pyrogenic fever in response to a pathogen (Lazarus et al., 2007). The MnPO projects to the dorsomedial hypothalamus (DMH) and the raphe pallidus (RPa) and this circuit is necessary for the initiation of heat production by brown adipose tissue (Machado et al., 2020). Prostaglandin receptors (EP3R) within the MnPO are critical for febrile response initiation (Machado et al., 2020). To date, the impact of unresolved early life pain on fever initiation and sickness behavior following pathogen exposure has not been explored.

Given that the nervous, immune, and endocrine systems undergo high levels of plasticity during the perinatal period, we hypothesized that exposure to unresolved pain during this critical developmental time point would disrupt normal development, resulting in alterations in the central immune response to a pathogen. Here we report that inflammatory pain on the day of birth potentiates the febrile response and cellular activation in multiple brain regions in male and female rats compared to controls.

## Materials and Methods

### Animals

Timed-pregnant Sprague Dawley rats (Charles River) were obtained on gestational day 14 (E14) and housed individually under 12:12-hour light:dark cycle with ad libitum access to food and water. On the day of birth (P0), pups were sexed by examination of anogenital distance. All litters were reared identically, weaned at P21, and housed in groups of 2-3 with same sex littermates. Male and female rats were used in all experiments. All experiments adhered to the guidelines of the Committee for Research and Ethical Issues of IASP and were approved by the Georgia State University Animal Care and Use Committee.

### Early Life Manipulations

Within 24 hours of birth, male and female rat pups were injected with 5µL of 1% carrageenan (CGN; Sigma, St. Louis MO) dissolved in sterile saline into the intraplantar surface of the right hind paw.

Control pups were handled in a similar manner and returned to their home cage (LaPrairie and Murphy, 2007). Intraplantar saline injections were not used due to induction of mild inflammation (LaPrairie and Murphy, 2007). All pups within a litter received the same treatment.

### Surgery

In adulthood (P60), Thermocron iButtons were implanted into the peritoneal cavity to monitor core body temperature. Rats were initially anesthetized with 5% isoflurane and maintained at 2-3% isoflurane during surgery. Following an abdominal incision, a wax coated iButton was inserted into the intraperitoneal cavity and the muscle wall closed with silk sutures. Carprofen (5 mg/kg/mL, sub- q) was administered preoperatively and 24 hours post-surgery for acute pain management.

### LPS Induced Immune Challenge

10-14 days following iButton implantation, rats were administered lipopolysaccharide (LPS, 250 µg/kg/mL; i.p.) to induce a fever (Konsman et al., 1999; Roth & Blatteis, 2014). Core body temperature was recorded in 10-minute intervals by the iButton loggers. Rats were assessed every 2- hours for sickness behaviors using an 8-point Likert scale based on changes in physical appearance (ear flattening, nose/cheek flattening, orbital tightening, and piloerection). Rats were sacrificed via decapitation at one of 3 time points post-LPS: 24 hours, peak fever (as determined by 24hr fever data), or 2 hours (fever initiation).

### Tissue Collection and Immunohistochemistry

Following decapitation, trunk blood was collected and the iButton recovered from the abdomen. Brains were extracted and drop fixed in a solution of 4% paraformaldehyde for 24 hours then stored in 30% sucrose solution at 4°C. Brains were sectioned coronally at 40µm on a Leica 2000R freezing microtome and stored free-floating in cryoprotectant-antifreeze solution at -20°C. Sections containing MnPO were selected and rinsed in KPBS. Sections were then incubated in primary antibody solution containing Rabbit anti-EP3R (1:500, Cayman Chem, 101760), Mouse anti-VGlut2 (1:1k, EMD Millipore, MAB5504), and Guinea Pig anti-VGat (1:3k, Synaptic Systems, 131004) overnight at 4°C. Following KPBS rinse, sections were incubated in a secondary antibody solution containing AF555 donkey anti-rabbit (1:200, ThermoFisher, A31572), AF488 donkey anti-mouse (1:200, Life Technologies, A21202), AF647 donkey anti-guinea pig (1:200, Jackson ImmunoResearch, 706-605-148), and DAPI (0.15uL/mL, Santa Cruz Biotech, sc-3598) for 2 hours. Sections were then washed in KPBS, mounted onto slides, and coverslipped with SlowFade Diamond fluorescent mounting media (Life Technologies, S36972), and then stored at 4°C in the dark.

Sections containing the organum vasculosum of the lamina terminalis (OVLT) were selected and processed as described above with primary antibody Rabbit anti-Cox-2 (1:1k, Cayman Chem, 160123) and secondary antibody AF555 donkey anti-rabbit (1:200, ThermoFisher, A31572).

### Fos Immunolabeling

Whole brain analysis of LPS induced Fos expression was conducted. A 1:6 series of brain slices were processed for Fos (1:30k, ab208942, Abcam) as described above. Sections were mounted onto gelatin subbed slides, air dried, counterstained with cresyl violet, and cleared with xylenes. Slides were then coverslipped with Permount mounting media.

### Imaging and Quantification

Fluorescent slides were imaged on a laser scanning confocal microscope (Zeiss, LSM 700) using the 40x Plan-Apochromat N.A. 1.4 with oil immersion objective. Z-stack images were displayed as maximum intensity projections of the entire z-series using Zen black software (Zeiss). Each image was analyzed for the total number of cells as well as the number of Fos+ cells (dual labeled with Ni- DAB and DAPI). Signal intensity for each fluorescent channel was taken as a measure of the expression level of its associated receptor protein.

Brightfield images of the Fos labeled slides were acquired on a Nikon Eclipse E800 microscope with a QImaging Retiga EXi CCD camera using the 4X Plan-Apochromat N.A. 0.2 and 10X Plan- Apochromat N.A. 0.45 objectives. Values analyzed represent the average number of Fos+ cells from two slices per region. Counts were performed using Fiji (NIH) or by hand by at least two people. All cell counts were conducted blind to experimental condition.

### Prostaglandin Quantification

Tissue for PGE2 analysis was frozen on dry ice and stored at -80°C. The anterior medial POA was dissected using a razor blade with cuts made at bregma -0.96mm and +1.08mm. Frozen tissue was homogenized on ice with a motorized pestle until no visible solid particles remain in ELISA extraction buffer comprised of 100 mM PIPES (pH 7), 500 mM NaCl, 0.2% Triton, 2 mM EDTA, 5 µg/mL Aprotinin, 0.1 µg/mL Pepstatin A, and 0.5 µg/mL Antipain. Samples were then centrifuged at 5,000 x g for 10 minutes at 4 °C. Supernatant was collected and protein concentration determined with Bradford Protein assay. All samples were diluted to 2 mg/mL with ELISA extraction buffer, aliquoted, then stored at -80 °C until ready to use. PGE2 levels were quantified via ELISA (ADI-901- 001, Enzo Biological).

### Statistical Analysis

Data are presented as mean ± SEM. All measurements and analyses were performed blind to treatment group. Data were assessed for normality and homogeneity of variance using Shapiro-Wilk and Bartlett’s tests. Points greater than +/- 2 standard deviations from the mean were removed. To control for circadian rhythm, LPS temperatures were time matched to baseline 24hrs prior, and the difference score calculated. Center of gravity analysis was conducted as a measurement of the mean phase of the curve, change in temperature weighted over time, and was reported as hours post-injection of LPS. This was calculated by taking the time average of the temperature change over the duration of each individual rat’s fever and calculating center of gravity as 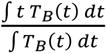 (*t* = time in hours, *T_B_* = core body temperature, *T_B_(t)* = core body temperature as a function of time; Paul et al., 2011; Kreyszig, 1993). Data were analyzed using t-test (k=2) or three-way or two-way mixed model analysis of variance (ANOVA; k>2) for each measure where appropriate. Temperature, sickness behavior, and brain region were treated as within subject factors; sex and P0 treatment were considered between subject factors. Post hoc comparisons were performed using Tukey’s HSD. All statistical tests were performed using GraphPad Prism (version 9.3.1) with α set at 0.05.

## Results

### Early life pain alters fever initiation and peak in females

To determine if ELP impacted the febrile response to a pathogen, rats were administered LPS and their body temperature recorded for 24 hours. Fever was defined as an increase of >0.75°C maintained for at least 3 consecutive hours. All rats showed a handling-induced spike in their temperature lasting 1 hour post-administration of LPS (Fig. 1A). All four groups showed an increase in body temperature beginning ∼3-5 hours post-LPS that lasted ∼5-7 hours (Fig. 1A). By 14hrs post-LPS, all rats showed recovery from fever, defined as 3 consecutive hours with <0.25°C difference from baseline (Fig. 1B). No effect of sex (F_(1,30)_=0.9451, p>0.05) or treatment (F_(1,30)_=0.06713, p>0.05) was noted for fever duration, and there was no significant interaction (F_(1,30)_=0.01967, p>0.05). As fever was limited to hours 0-14 in all treatment groups, only these hours were included for further analysis. Three-way ANOVA revealed significant Time*Sex (F_(89, 2670)_=1.327; p=0.0234) and Time*Sex*Treatment interactions (F_(89, 2670)_=1.289; p=0.0375) for temperature change during hours 0-14. Post-hoc analysis revealed that overall, LPS induced a significantly higher febrile response in ELP females compared to control females that is most evident in hour 8 in which the average increase in body temperature in ELP females was 1.600±0.052°C compared to control females at 1.063±0.019°C (Fig. 1C; F_(1,16)_=4.619; p=0.0473). No significant effect of treatment was seen in males (Fig. 1E; F_(1,17)_=1.158; p>0,05). Overall peak temperature was also significantly greater in ELP females with an increase of 0.353 ± 0.174°C compared to control females (Fig. 1C; t_(14)_=2.033; p=0.0307). No effect of early life treatment was seen in males. Center of gravity analysis, used to assess for shift in the febrile response as a function of time, also revealed a significant Treatment*Sex interaction (F_(1,28)_=10.79, p=0.0027). Post-hoc analysis revealed a significant rightward phase shift in ELP females (Fig. 1D; t_(28)_=2.704; p=0.0229). The center of gravity for handled females’ temperature curve was 7.135 hours post-LPS compared to 8.238 hours post-LPS in ELP females, overall a positive 1.103 ± 0.3378 hour shift. No effect of treatment was seen in male groups (Fig. 1F; t_(28)_=1.940; p>0.05). Together, these analyses show that ELP females exhibit both an augmented fever and a delayed phase shift in their fever response compared to handled controls; no effect of early life treatment on febrile response to LPS was observed in males.

**Figure 1.**
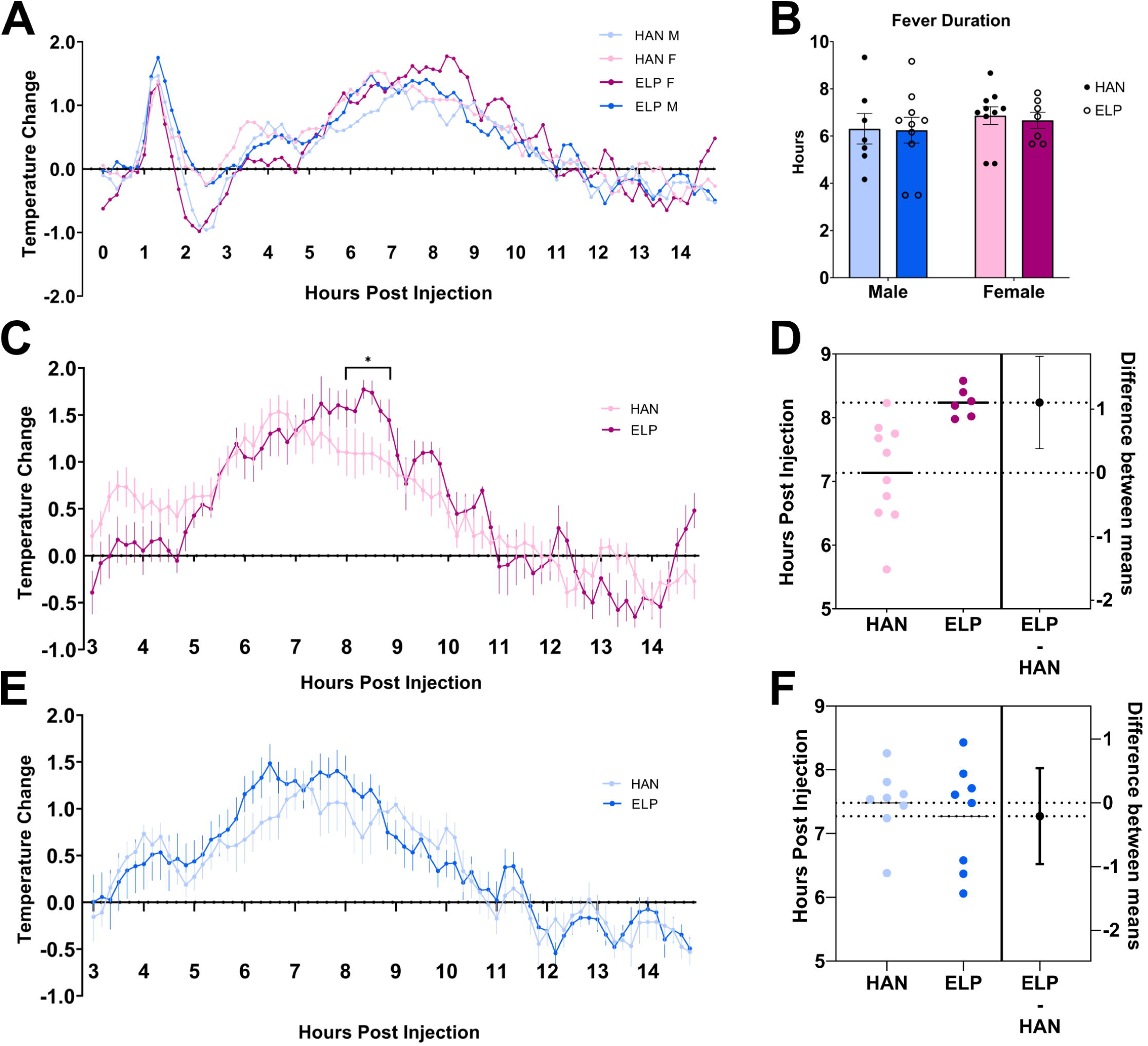
ELP results in increased fever severity compared to handled controls. Temperature change as a function of time in male and female rats with the time of LPS injection indicated by the red arrow (A). No differences were seen in the overall duration of fever (B). Female rats exposed to ELP show an increased change in peak temperature (p=0.0307) and are significantly higher at hour 8 (p=0.0473) (C) and a rightward shift in fever response measured as a shift in the center of gravity (D, p=0.009). Male rats exposed to ELP show a trend toward increased change in temperature (E) but no shift in fever response (F, p=0.1127).

### Early life pain increases female sickness behaviors

We next evaluated the impact of ELP on LPS induced sickness behaviors. Sickness behavior was evaluated every 2 hours using an 8-point Likert scale for 4 behaviors: ears flattening against the head, nose/cheek flattening, orbital tightening, and fur piloerection (Fig. 2A). Sickness score as a function of time is shown from 2 hours post injection through hour 16 when all rats no longer showed any sickness behaviors (Fig. 2B). Two-way ANOVA of peak sickness behavior revealed a Treatment*Sex interaction (Fig. 2C; F_(1,23)_=4.596; p=0.0428). Post-hoc analysis revealed a sex difference between handled groups (p=0.0014) and a treatment effect in females only (Fig. 2C; p=0.0068). Overall, sickness behavior increased in females as a function of treatment. No similar changes were observed in males. Correlation of peak sickness score to peak temperature indicates that changes in sickness behavior are associated with increased temperature change (Fig. 2D; R^2^=0.5512, p=0.0002).

**Figure 2.**
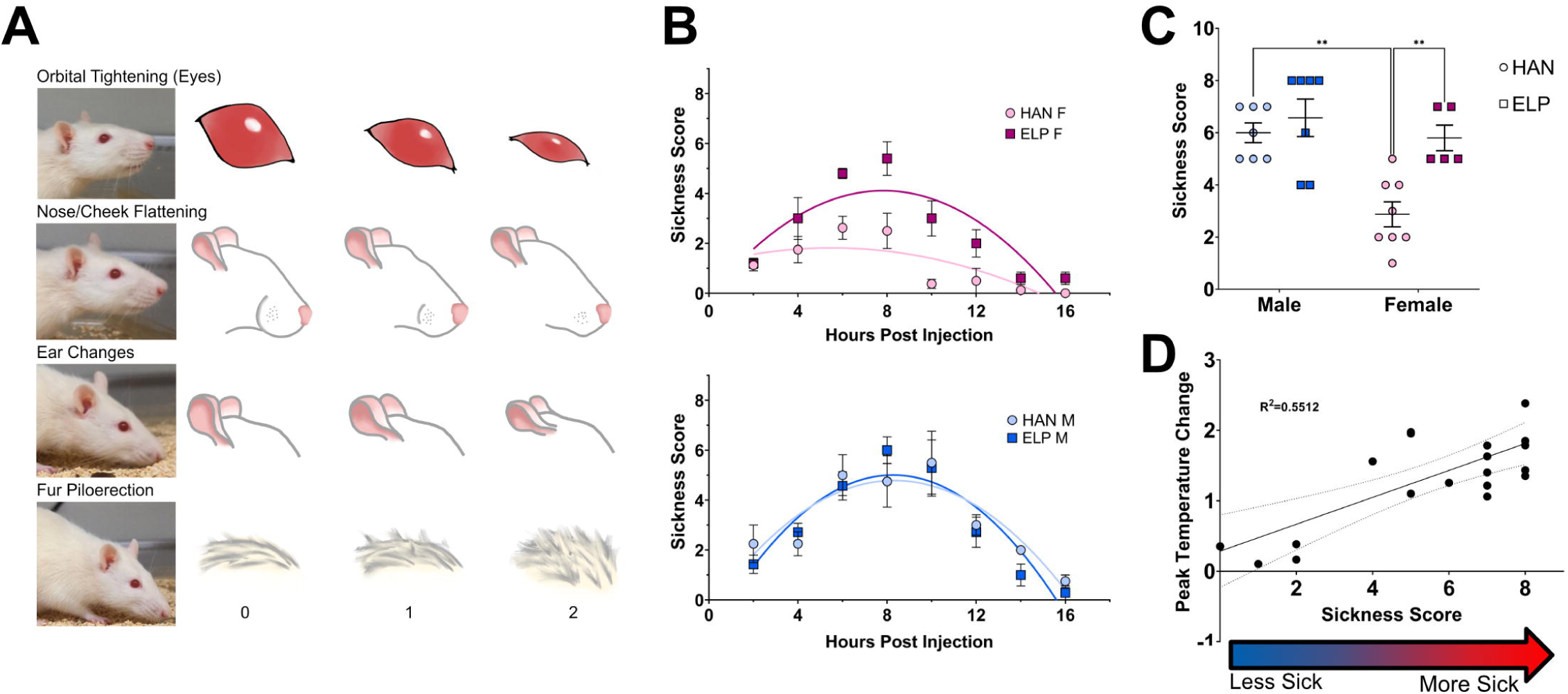
ELP results in increased sickness behaviors in females but not males. Sickness behaviors were quantified as shown with each behavior receiving a score of 0-2 and a composite score for each animal was totaled out of 8 (A). The change in sickness score was measured every 2 hours to produce a sickness curve. Parabolic regression lines were fitted for visualization (B). Peak sickness score showed increase in severity in ELP females compared to handled females (C, p=0.0053). A sex difference was also seen between our handled groups (p=0.0029). Males showed no differences between treatment groups. Peak temperature change and peak sickness score are strongly correlated (D, R^2^=0.5512, p=0.0002) in all groups except for handled females as they show little to no signs of sickness during the fever.

### Early life pain rats show altered levels of cellular activation during fever initiation

Thus far, our studies have shown that ELP results in a significant increase in febrile response and sickness behavior in female, but not male, rats. To elucidate the potential neural circuitry driving the observed differences in fever and sickness behavior, we next examined brain activation patterns during fever. To assess what may be driving the differences observed in fever initiation, whole brain analysis of Fos expression was conducted at 2 hours post LPS injection. Three-way ANOVA of Fos+ cell counts showed an overall interaction between Region*Treatment*Sex (Fig. 3; F_(13,126)_=4.373, p=0.0005). Subsequent two-way ANOVAs showed an overall effect of treatment in the males (Fig. 3A; F_(1,12)_=19.85, p=0.0008) with ELP males showing a significantly lower level of Fos expression. Post-hoc analysis for individual regions revealed a significant decrease in ELP males in the NAcSh (p=0.0008), the MnPO (p=0.0076), and the MeA (p=0.0101) compared to handled controls. The MnPO is critical for driving the inflammatory fever response, while the NAcSh and MeA are both associated with stress pathways, likely related to the observed sickness behaviors. Two-way ANOVA of the female groups showed a significant effect of treatment (Fig. 3A; F_(1,9)_=9.076, p=0.0147) with ELP females having increased levels of Fos expression compared to their handled counterparts. Post-hoc analysis revealed no effect of ELP treatment within individual regions in females. Additionally, correlational analysis of Fos expression between brain regions was conducted for each group to assess whether global patterns of Fos expression were impacted by ELP (Fig. 3B). ELP rats of both sexes showed increases in the number of significant region correlations compared to handled rats.

**Figure 3.**
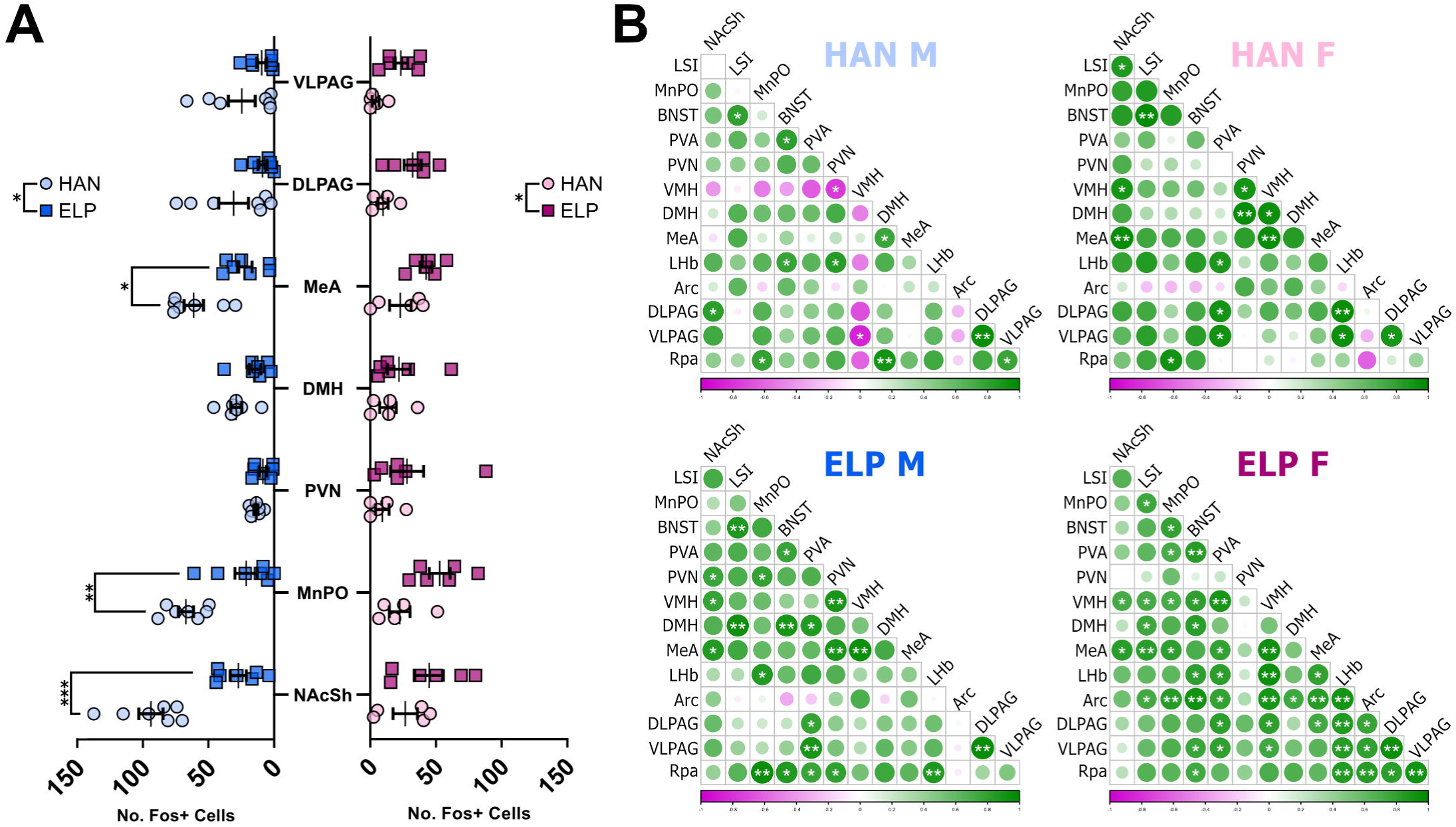
ELP alters cellular activation during fever initiation. Whole brain analysis of Fos expression during fever initiation showed differential changes in ELP males and females. ELP males showed an overall decrease in Fos expression compared to handled males (p=0.0008) with the NAcSh (p=0.0008), MnPO (p=0.0076), and the MeA (p=0.0101) showing significantly lower expression (A). ELP females showed an overall increase in Fos expression compared to handled females (p=0.0147) with no differences seen in individual regions (A). Correlational analysis between regions for each group showed an increase in the number of correlated regions in both male and female ELP rats (B).

### Early life pain rats show increased levels of cellular activation at fever peak

To assess what may be driving the differences observed at peak fever, analysis of whole brain Fos expression was conducted at each group’s respective peak (6 hours post LPS for HAN males and females and ELP males; 8 hours post LPS for ELP females). Two-way ANOVA of the number of brain regions showing LPS induced Fos revealed an increase as a main effect of ELP treatment (Fig. 4A; F_(1,19)_=6.54, p=0.0193). Post-hoc analysis showed this effect was driven by ELP females compared to handled controls (Fig. 4A, p=0.0482). Three-way ANOVA of Fos+ cell counts showed an overall main effect of Treatment in which ELP rats had elevated numbers of Fos+ cells compared to handled controls (F_(1,18)_=11.63, p=0.0031) as well as a Region*Treatment (F_(6,108)_=2.261, p=0.0428) and a Region*Sex (F_(6,108)_=3.960, p=0.0013) interaction. Subsequent two-way ANOVAs to determine what is driving this increase showed a main effect of Treatment (F_(1,8)_=41.16; p=0.0002) and a Region*Treatment interaction in females (F_(6,48)_=9.582; p<0.0001) with ELP females showing a higher number of Fos+ cells (Fig. 4B). No effect was seen in the male groups (Fig. 4B). Post-hoc analysis from two-way ANOVAs also revealed sex differences where handled females showed lower Fos expression than handled males in the MnPO (p=0.022) and PVA (p=0.0015). The PVN also showed decreased expression in ELP females compared to ELP males (p=0.0014). Together these suggest that ELP females show more activated regions and more activation within those regions than their handled counterparts at peak. In contrast, no differences were noted for ELP vs. HAN males in Fos expression at peak fever.

**Figure 4.**
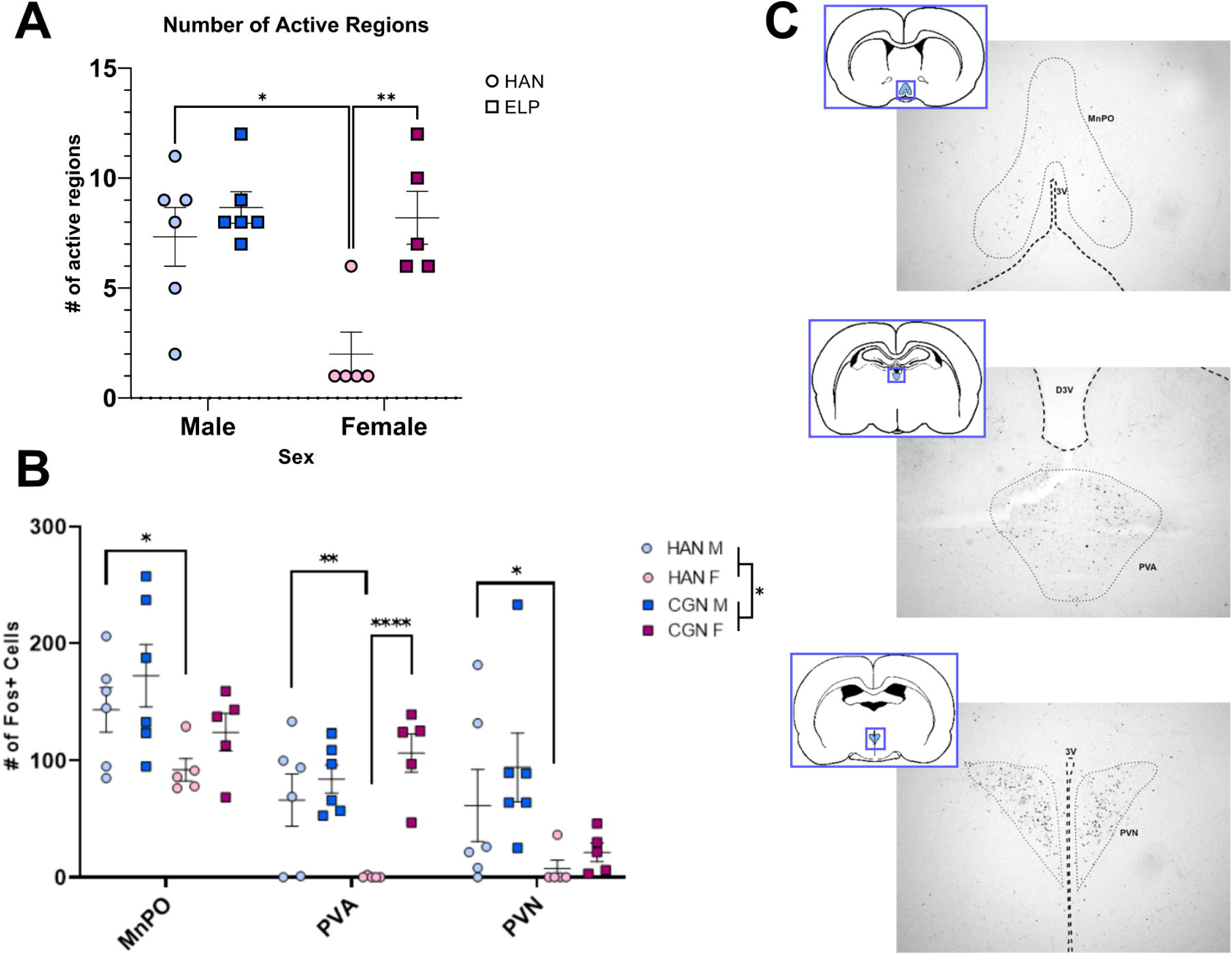
ELP results in increased c-Fos expression at peak fever in regions associated with fever and sickness responses. Whole brain analysis of Fos expression at fever peak showed an overall increase in the number of active regions in ELP rats (A) with an increase in ELP females compared to handled controls (p=0.0482). Additionally, ELP rats showed an increase in the number of Fos+ cells within the highest expressing regions. Handled females showed significantly lower expression overall compared to handled males, particularly in the MnPO (B; p=0.022), PVA (p=0.0015), and the PVN (p=0.292). Within the PVA ELP females also show higher expression of Fos+ cells compared to handled females (p=0.0044). Representative images show the MnPO, PVA, and PVN regions of interest (C).

### Early life pain rats show increased PGE2 and Cox-2 at 2 hrs post LPS

Prostaglandin E2 is critical for the generation of a fever in response to a pathogen. In response to peripheral release of cytokines, Cox-2 is expressed to synthesize PGE2 which then acts on the EP3 receptor in the MnPO. In order to determine if differences in LPS induced febrile response were due to a difference in PGE2 production, we next examined the levels of PGE2 in the anterior medial preoptic area. Brain lysates from rats sacrificed either at 2 hours post LPS or at peak fever were analyzed for PGE2 levels (Fig. 5A-B). At 2 hours post LPS, no sex differences were detected, so data were collapsed to increase power. ELP rats showed an increase in PGE2 levels at 2 hours post LPS (Fig. 5A; t_(7.944)_=2.285; p=0.0519). At peak fever, PGE2 levels are shown as a fold change compared to saline controls (Fig. 5B). Two-way ANOVA was used to assess for significant main effects of sex and early life treatment. Analysis revealed a significant Sex*Treatment interaction (F_(1,12)_=5.679; p=0.0346) with no significant main effects of sex or treatment. Post-hoc analysis revealed a non-significant increase in ELP males compared to handled males (p=0.0978) and no differences in female groups.

**Figure 5.**
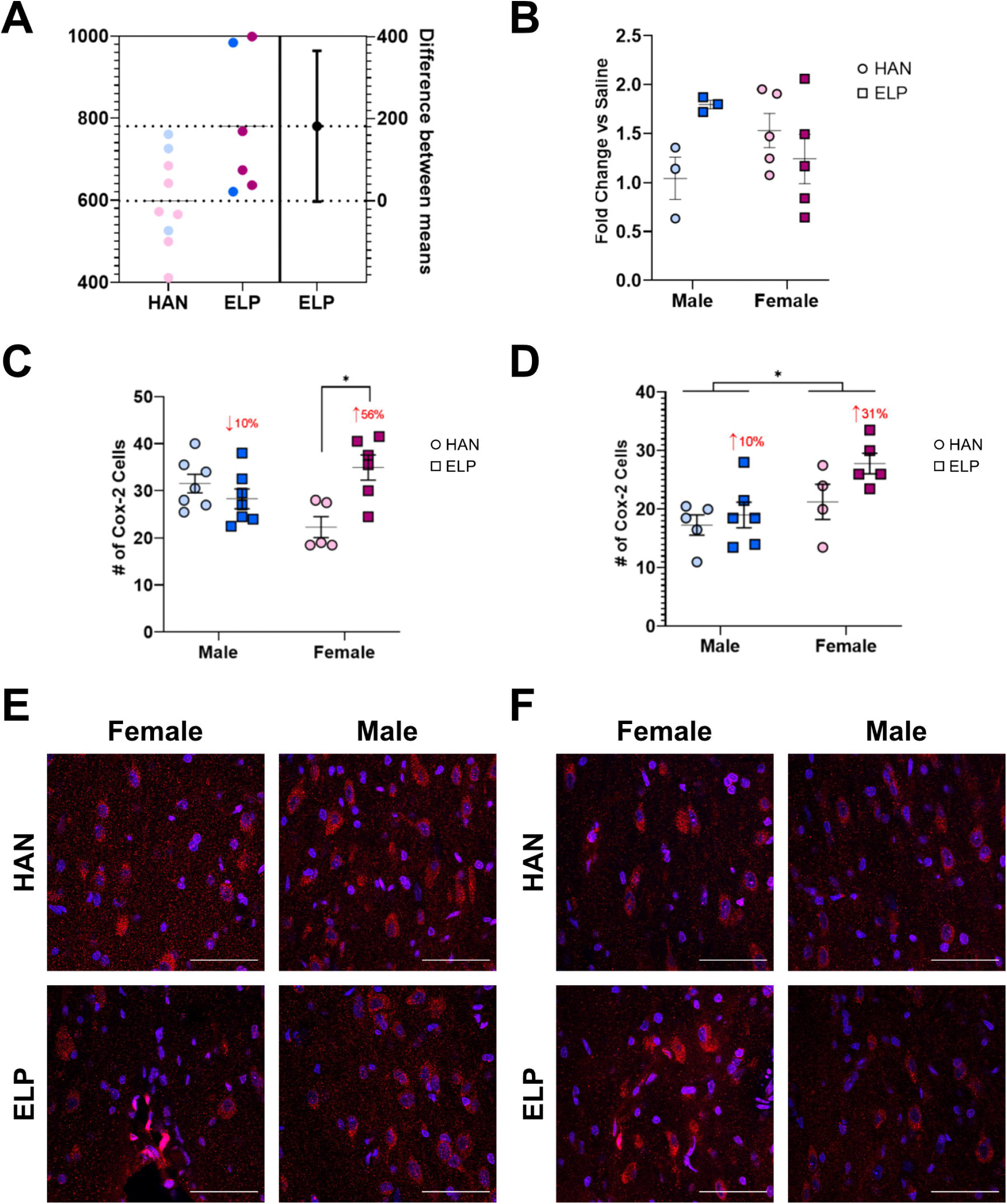
Early life pain rats show increased PGE2 and Cox-2 at 2 hrs post LPS. Brain POA homogenates were processed via ELISA for PGE2 levels (A-B). Analysis of PGE2 levels (pg/mL) at 2 hours post LPS revealed an increase in PGE2 in ELP rats (A, p=0.0519). At peak fever, PGE2 levels are shown as fold change compared to saline injected controls (B). Analysis yielded a significant Sex*Treatment interaction (p=0.0346) and post-hocs showed a non-significant increase in ELP males compared to handled males (p=0.0978) and no differences in females. Tissue containing OVLT was stained for Cox-2 and counterstained with DAPI (C-F). Analysis of Cox-2 positive cells at 2 hours post LPS (C) revealed a significant Sex*Treatment interaction (p=0.0021) and post-hocs showed higher expression in handled males compared to handled females (p=0.0482) and higher expression in ELP females compared to handled females (p=0.0064). Analysis of Cox-2 at peak fever (D) revealed a main effect of sex with females having more positive cells compared to males (p=0.0100) and a trend toward a main effect of ELP treatment (p=0.0769). Representative images of Cox-2 expression at 2 hours post-LPS (E) and peak fever (F). Scale bar = 50µm.

Cox-2 expression in the OVLT was assessed via IHC and quantified by the number of cells that were double positive for DAPI and Cox-2. Values for each rat were determined by averaging the number of positive cells from two slices (Fig. 5C-F). At 2 hours post LPS, analysis showed a significant Sex*Treatment interaction (F_(1,21)_=12.24; p=0.0021); no significant main effects of treatment or sex were found. Post-hoc analysis revealed a significant increase in handled males compared to handled females (p=0.0482) and a significant increase in ELP females compared to handled females (p=0.0064). At peak fever, analysis showed a significant main effect of sex (F(_1,16)_=8.542; p=0.0100) and a trend toward a main effect of treatment (F_(1,16)_=3.576; p=0.0769) with no interaction. Post-hoc analysis did not reveal any individual group differences. Together these results show that in response to LPS challenge ELP rats, particularly females, show elevated expression of the Cox-2 enzyme and subsequent elevation of PGE2 at 2 hours post-LPS. At peak fever, Cox-2 expression is elevated in females compared to males with a slight elevation in ELP rats compared to handled rats and no significant differences in PGE2 levels.

### Early life pain alters LPS induced receptor density in the MnPO

The MnPO is a critical region for the initiation of fever in response to a pathogen; thus we next examined if the differences observed in LPS induced febrile response were associated with alterations in GABA/Glutamate signaling in the MnPO. The role of the PGE2 receptor EP3R was also examined. Brain tissue from rats sacrificed either for basal levels (no LPS) or at peak fever was analyzed for levels of EP3R, VGlut2, and VGat (Fig. 6). To normalize receptor expression across individual subjects, expression values are reported as signal intensity/DAPI for each individual.

**Figure 6.**
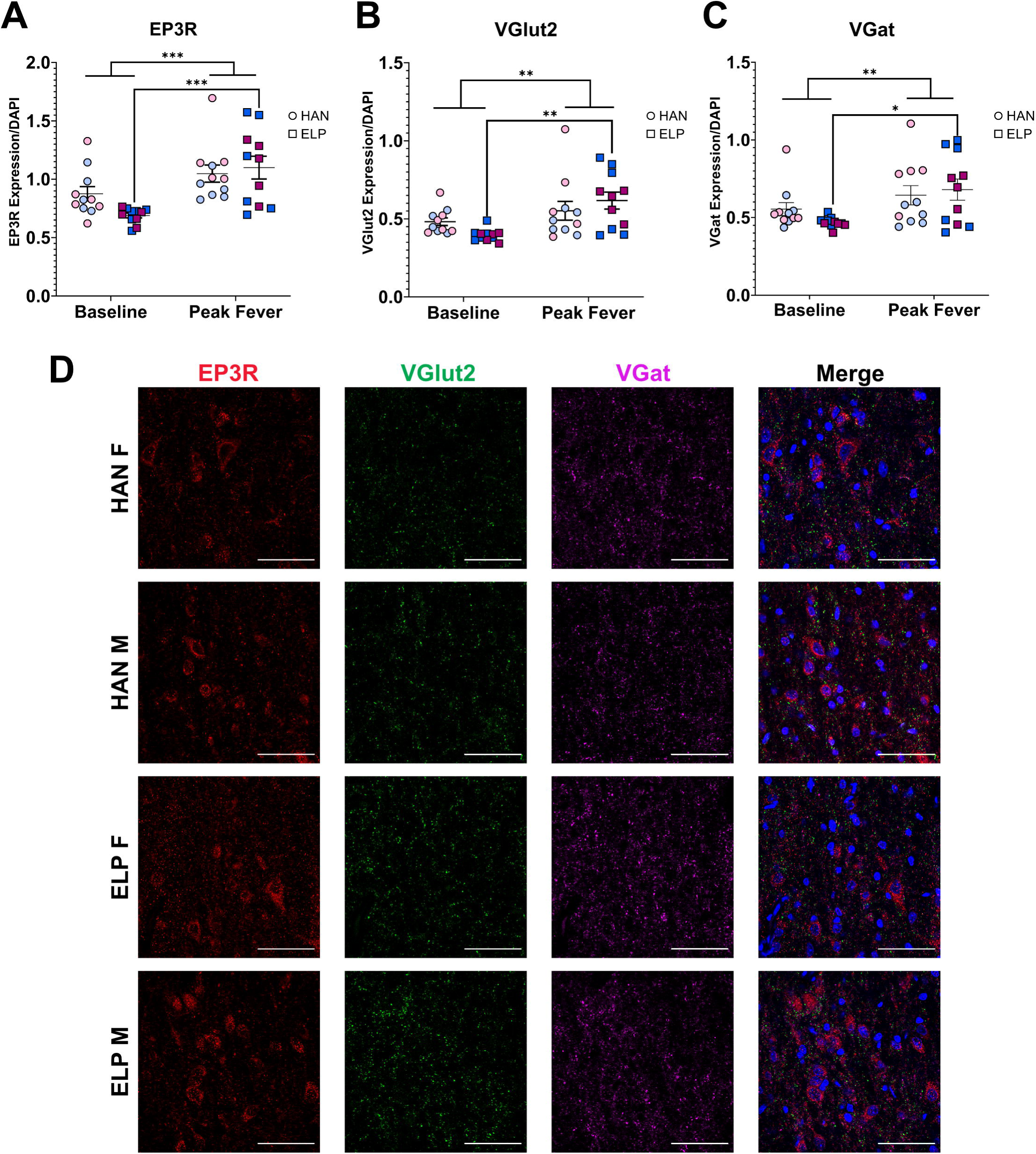
Early life pain alters LPS induced EP3 receptor and and GABA and Glutamate vesicular transporter density in the MnPO. MnPO tissue was stained for prostaglandin receptor EP3R (red), Vesicular glutamate transporter 2 (VGlut2, green), vesicular GABA transporter (VGat, magenta), and counterstained with DAPI (blue). Analysis of signal intensity/DAPI yielded no differences between groups in the expression of EP3R (A), VGlut2 (B), and VGat (C) at baseline or peak fever. Groups are shown collapsed across sex with individual points (pink=female, blue=male). There was a main effect of fever and all groups showed elevated expression in all signals at peak fever compared to baseline. ELP rats in particular showed a significant increase in EP3R (A, p=0.0008), VGlut2 (B, p=0.0038), and VGat (C, p=0.0271) at peak fever compared to baseline. Representative images of MnPO expression at peak fever (D, scale bar = 50µm).

Three-way ANOVA was used to assess for significant main effects of sex, treatment, and fever. There was no significant main effect of sex so these data are collapsed for data presentation. No impact of ELP was noted for EP3R (Fig. 6A; F_(1,40)_=0.96, p>0.05), VGlut2 (Fig. 6B; F_(1,40)_=0.06, p>0.05), or VGat (Fig. 6C; F_(1,40)_=0.24, p>0.05). In contrast, LPS induced a significant increase in all three signals at peak fever compared to basal expression levels (EP3R: F_(1,40)_=17.84, p=0.0001; VGlut2: F_(1, 40)_=11.63, p=0.0015; VGat: F_(1,40)_=8.79, p=0.0052). Post-hoc analysis revealed that this increase in all three signals was driven by ELP treatment (EP3R: p=0.0008; VGlut2: p=0.0038; VGat: p=0.0271). Together these results suggest that LPS-induced fever results in a significant increase in signal expression that is augmented in ELP rats, independent of sex.

## Discussion

The present studies are the first to show the long-term effects of unresolved early life pain on adult immune function. Here we show that early life pain in females resulted in a stronger central response to LPS, observed as a higher fever and increased severity of sickness behaviors.

### Early life pain alters febrile response and sickness behaviors in females

Male and female rats that experienced unresolved early life pain had a higher peak fever response than their handled counterparts. This was particularly evident in the female groups which had a significant increase in peak temperature in the ELP rats compared to handled rats. Fever generation is critical for a successful defense against a pathogen, however too high of a deviation from normal body temperature can result in additional damage to major organ systems. For this reason, body temperature and the response to a pathogen are closely controlled via communication between peripheral immune cells and the brain. In addition to an increase in their peak fever, ELP females also had a delayed response in their fever time-course. In comparison, the handled control females mount a fever response within 3 hours of LPS administration but show a lower peak temperature change. ELP females also showed an increased severity of sickness behaviors compared to handled females, which notably did not exhibit many sickness behaviors. Male rats exposed to early life pain also show an increase in their peak temperature change, but to a smaller degree than in the female groups. No shift was noted in the time course of their fever or in their sickness behavior, though this may be due to a ceiling effect as all males scored high on sickness behavior severity.

### Early life pain alters LPS induced Fos activation at fever initiation and peak

This delay is not likely due to a lack of brain activation as our ELP females show greater activation at 2 hours post-LPS compared to their handled counterparts. Rather, this suggests that communication between the peripheral immune system and central neural circuits underlying fever generation have been altered by ELP. Handled female rats also show lower levels of Fos activation during fever initiation and at fever peak, suggesting that their peripheral immune system could be more efficiently responding to the immune challenge. This agrees with the clinical data in which females show greater innate immune responses, but their adaptive immunity is prone to over-reacting, resulting in autoimmune responses (Lasrado et. al, 2020; Klein et al., 2015; Taneja, 2018). Priming of this system by ELP could drive the immune response more toward a hyperresponsive state, resulting in the increased levels of Fos activation seen in ELP females. In contrast, ELP males show overall lower levels of Fos expression during fever initiation but elevated levels at peak fever compared to their handled counterparts. Further studies looking into the potential priming effects of ELP in a sex specific manner on the immune response during an LPS challenge are needed to better understand the changes we see here.

### Early life pain rats show increased PGE2 and Cox-2 at 2 hrs post LPS

PGE2 is critical for the generation of a fever during an immune challenge through it’s actions in the MnPO. Synthesis of PGE2 is dependent on the action of Cox-2 which is inducible in response to inflammation, such as cytokines. Since PGE2 cannot readily cross the blood brain barrier, this synthesis must occur locally and the OVLT is situated adjacent to the MnPO. At 2 hours post-LPS during fever initiation ELP rats of both sexes show increased PGE2 in the preoptic area of the brain, but only ELP females show increased expression of Cox-2 in comparison to handled females. This follows the same pattern as seen with Fos expression in ELP rats during fever initiation. At peak fever there are no significant differences in preoptic PGE2 levels and a smaller increase in Cox-2 expression with females having more Cox-2 positive cells compared to males. These results further suggest a priming effect of ELP that results in a greater inflammatory response during an immune challenge that in females may be working in the early phase of the response to LPS.

### Early life pain augments LPS induced MnPO signaling increases

Signaling in the MnPO is critical for the generation of a pyrogenic fever in response to a pathogen. Alterations in MnPO signaling are capable of driving changes in the fever response through GABA/Glutamate signaling and prostaglandin receptor activation. LPS administration resulted in upregulation of both VGlut2 and VGat expression as well as EP3R compared to basal expression levels in all rats. No impacts of sex were noted, but ELP treatment resulted in an augmented increase in expression levels compared to handled controls. This increase parallels the elevated fever seen in ELP rats but does not explain the greater difference seen in ELP females compared to males.

In conclusion, our model to assess the long-term effects of ELP on immune function shows that females are more vulnerable to the immune consequences of ELP with their increased fever and sickness severity. ELP females also have greater levels of cellular activation across multiple brain regions, many of which are associated with febrile and stress response. ELP males do not show any significant differences in fever or sickness behaviors but have altered PGE2 levels, MnPO signaling and global cellular activation responses that warrant further investigation into other parts of the immune response such as peripheral immune signalling or microglial response within the MnPO to understand how ELP is impacting male immune responses. More specific methods of assessing sickness behavior responses may also help to determine if any changes are occurring in the male groups which were not able to be captured in this study.

## Author Contributions

Morgan Gomez: Conceived and performed all experiments, quantified and analysed all data, and wrote and edited the manuscript.

Melani Macik: Assisted with performing of the experiments and data quantification and analysis

Hannah Harder: Assisted with experimental design, performing of the experiments, data analysis and edited the manuscript.

Malika Pandit: Assisted with performing of the experiments and data quantification and analysis

Anne Murphy: Conceived the experiments, wrote and edited the manuscript, and supervised the study.

## Acknowledgments

We would like to acknowledge Lauren Hanus for help in training on the ELP protocols, surgeries, and immunohistological techniques; Chris Searles for assisting on surgeries and behavioral protocols; and Claudia Sanabria for training on confocal microscopy techniques.

## References

Berrington, J. E., Barge, D., Fenton, A. C., Cant, A. J., & Spickett, G. P. (2005). Lymphocyte subsets in term and significantly preterm UK infants in the first year of life analysed by single platform flow cytometry. Clinical and Experimental Immunology, 140(2), 289–292. 10.1111/j.1365-2249.2005.02767.x

Carbajal, R., Rousset, A., Danan, C., Coquery, S., Nolent, P., Ducrocq, S., Saizou, C., Lapillonne, A., Granier, M., Durand, P., Lenclen, R., Coursol, A., Hubert, P., Blanquat, L. de S., Boëlle, P.- Y., Annequin, D., Cimerman, P., Anand, K. J. S., & Breart, G. (2008). Epidemiology and Treatment of Painful Procedures in Neonates in Intensive Care Units. 11.

Dourson, A. J., Ford, Z. K., Green, K. J., McCrossan, C. E., Hofmann, M. C., Hudgins, R. C., & Jankowski, M. P. (2021). Early Life Nociception is Influenced by Peripheral Growth Hormone Signaling. The Journal of Neuroscience, 41(20), 4410–4427. 10.1523/JNEUROSCI.3081-20.2021

Grunau, R. E., Cepeda, I. L., Chau, C. M. Y., Brummelte, S., Weinberg, J., Lavoie, P. M., Ladd, M., Hirschfeld, A. F., Russell, E., Koren, G., Van Uum, S., Brant, R., & Turvey, S. E. (2013).

Neonatal Pain-Related Stress and NFKBIA Genotype Are Associated with Altered Cortisol Levels in Preterm Boys at School Age. PLoS ONE, 8(9), e73926. 10.1371/journal.pone.0073926

Klein, S. L., Marriott, I., & Fish, E. N. (2015). Sex-based differences in immune function and responses to vaccination. Transactions of the Royal Society of Tropical Medicine and Hygiene, 109(1), 9–15. 10.1093/trstmh/tru167

Konsman, J. P., Kelley, K., & Dantzer, R. (1999). Temporal and spatial relationships between lipopolysaccharide-induced expression of fos, interleukin-1 β and inducible nitric oxide synthase in rat brain. Neuroscience, 89(2), 535–548. 10.1016/S0306-4522(98)00368-6

Kreyszig, E. (1993). Advanced Engineering Mathematics (10th ed.). Wiley.

LaPrairie, J. L., & Murphy, A. Z. (2007). Female rats are more vulnerable to the long-term consequences of neonatal inflammatory injury. Pain, 132(Supplement 1), S124–S133. 10.1016/j.pain.2007.08.010

Lasrado, N., Jia, T., Massilamany, C., Franco, R., Illes, Z., & Reddy, J. (2020). Mechanisms of sex hormones in autoimmunity: Focus on EAE. Biology of Sex Differences, 11(1), 50. 10.1186/s13293-020-00325-4

Lazarus, M., Yoshida, K., Coppari, R., Bass, C. E., Mochizuki, T., Lowell, B. B., & Saper, C. B. (2007). EP3 prostaglandin receptors in the median preoptic nucleus are critical for fever responses. Nature Neuroscience, 10(9), 1131–1133. 10.1038/nn1949

Machado, N. L. S., Bandaru, S. S., Abbott, S. B. G., & Saper, C. B. (2020). EP3R-Expressing Glutamatergic Preoptic Neurons Mediate Inflammatory Fever. The Journal of Neuroscience, 40(12), 2573–2588. 10.1523/JNEUROSCI.2887-19.2020

March of Dimes, PMNCH, Save the Children, & WHO. (2012). Born too Soon: The Global Action Report on Preterm Birth (C. Howson, M. Kinney, & J. Lawn, Eds.).

Melville, J. M., & Moss, T. J. M. (2013). The immune consequences of preterm birth. Frontiers in Neuroscience, 7, 79. 10.3389/fnins.2013.00079

Osterman, M., Hamilton, B., Martin, J., Driscoll, A., & Valenzuela, C. (2021). Births: Final Data for 2020. National Center for Health Statistics (U.S.). 10.15620/cdc:112078

Paul, M. J., Indic, P., & Schwartz, W. J. (2011). A role for the habenula in the regulation of locomotor activity cycles: Habenula and activity cycles. European Journal of Neuroscience, 34(3), 478–488. 10.1111/j.1460-9568.2011.07762.x

Pelkonen, A. S., Suomalainen, H., Hallman, M., & Turpeinen, M. (1999). Peripheral blood lymphocyte subpopulations in schoolchildren born very preterm. Archives of Disease in Childhood - Fetal and Neonatal Edition, 81(3), F188–F193. 10.1136/fn.81.3.F188

Rodkey, E. N., & Pillai Riddell, R. (2013). The Infancy of Infant Pain Research: The Experimental Origins of Infant Pain Denial. The Journal of Pain, 14(4), 338–350. 10.1016/j.jpain.2012.12.017

Roofthooft, D. W. E., Simons, S. H. P., Anand, K. J. S., Tibboel, D., & van Dijk, M. (2014). Eight Years Later, Are We Still Hurting Newborn Infants? Neonatology, 105(3), 218–226. 10.1159/000357207

Roth, J., & Blatteis, C. M. (2014). Mechanisms of fever production and lysis: Lessons from experimental LPS fever. Comprehensive Physiology, 4(4), 1563–1604. 10.1002/cphy.c130033

Schwaller, F., & Fitzgerald, M. (2014). The consequences of pain in early life: Injury-induced plasticity in developing pain pathways. European Journal of Neuroscience, 39(3), 344–352. 10.1111/ejn.12414

Simons, S. H. P., van Dijk, M., Anand, K. S., Roofthooft, D., van Lingen, R. A., & Tibboel, D. (2003). Do We Still Hurt Newborn Babies?: A Prospective Study of Procedural Pain and Analgesia in Neonates. Archives of Pediatrics & Adolescent Medicine, 157(11), 1058. 10.1001/archpedi.157.11.1058

Strunk, T., Currie, A., Richmond, P., Simmer, K., & Burgner, D. (2011). Innate immunity in human newborn infants: Prematurity means more than immaturity. The Journal of Maternal-Fetal & Neonatal Medicine, 24(1), 25–31. 10.3109/14767058.2010.482605

Taneja, V. (2018). Sex Hormones Determine Immune Response. Frontiers in Immunology, 9, 1931. 10.3389/fimmu.2018.01931

Victoria, N. C., Inoue, K., Young, L. J., & Murphy, A. Z. (2013). A Single Neonatal Injury Induces Life-Long Deficits in Response to Stress. Developmental Neuroscience, 35(4), 326–337. 10.1159/000351121

Victoria, N. C., Karom, M. C., Eichenbaum, H., & Murphy, A. Z. (2013). Neonatal injury rapidly alters markers of pain and stress in rat pups. Developmental Neurobiology, 74(1), 42–51. 10.1002/dneu.22129

Zouikr, I., Bartholomeusz, M. D., & Hodgson, D. M. (2016). Early life programming of pain: Focus on neuroimmune to endocrine communication. Journal of Translational Medicine, 14(1), 123. 10.1186/s12967-016-0879-8

